# In-cell structural insight into the asymmetric assembly of central apparatus in mammalian sperm axoneme

**DOI:** 10.1101/2024.08.06.606614

**Authors:** Yun Zhu, Guoliang Yin, Linhua Tai, Binbin Wang, Yi Lu, Zhiyong Zhang, Suren Chen, Fei Sun

## Abstract

In motile cilia or flagella, the axoneme typically exhibits a “9+2” configuration, with the central apparatus (CA) consisting of extensively modified microtubules C1 and C2 to regulate ciliary motility. How C1 and C2 are interconnected asymmetrically remains unknown due to the lack of complete structural model of CA. Here, we utilized the *in situ* cryo-electron tomography approach to solve the in-cell structure of intact mouse sperm CA at sub-nanometer resolution and built its near-complete atomic model with the aid of AlphaFold2. We identified 39 different CA-associated proteins with 8 never reported from isolated specimen. We assigned the long chain-like ASH-containing proteins CFAP47 and HYDIN responsible for connecting C1 and C2. Sperm from *Cfap47*-knockout mice displayed a hollowing bridge in the CA structure, correlating with its reduced progressive motility. Our findings elucidate the molecular mechanisms of CA components in ciliary motility, and the implications of their mutations in human ciliopathies.

## Introduction

Cilia are evolutionarily conserved organelles that are present across most eukaryotic phyla. They are categorized into motile and non-motile subtypes, both sharing a common cytoskeletal scaffold known as the axoneme. Motile cilia facilitate cell movement and the transport of extracellular materials through rhythmic beats. In humans, cilia play crucial roles in various life processes, such as embryonic development, fertility, airway function, and cerebrospinal fluid circulation^1^. Defects in ciliary structures and functions can result in ciliopathies, a group of diseases such as congenital heart defects, hydrocephalus, and primary ciliary dyskinesia (PCD)^2,3^.

The axoneme can adopt either the “9+2” pattern or “9+0” pattern^4,5^. In motile cilia or flagella, the axoneme typically displays a “9+2” pattern, comprising 9 peripheral microtubule doublets (DMTs), inner and outer dynein arm (IDA and ODA), radial spoke (RS), and nexin-dynein regulatory complex (N-DRC), surrounding a pair of extensively modified microtubules named the central apparatus (CA) ^6,7^. While there are rare exceptions^8,9^, the CA primarily regulates ciliary motility by serving as a mechanical-force distributor that interacts with RSs to convey mechanochemical signals throughout the axoneme^10–12^. Cilia and flagella from various organisms, such as protists^13^, echinoderms^14^, and mammals^15^, have been utilized to investigate the CA. However, due to the dynamic nature and numerous components of the CA, research on its structure has been limited to overall morphology and component characterization for an extended period^16^. Generally, the CA exhibits a conserved basic structure, where two singlet microtubules, referred to as C1 and C2, are connected by structures collectively known as the bridge. Multiple projections are attached to each microtubule, extending towards the RSs. The C2 projections and bridge exhibit repeats with a 16 nm periodicity, while the C1 projections demonstrate both 16 nm and 32 nm periodicities^16,17^.

Two years ago, the high-resolution CA structure was determined using cryo-electron microscope (cryo-EM) with purified sample from *Chlamydomonas reinhardtii* (*C. reinhardtii*), providing insights into the structures of dissociated C1 and C2 microtubules, and their role in regulating ciliary beating^18,19^. The CA consists of six projections on the C1 microtubule (C1a–f) and five on the C2 microtubule (C2a–e). Functionally related projection proteins of the CA are clustered on a spring-shaped scaffold protein PF16, with some projection complexes containing a rachis-like protein that organizes all other subunits on the scaffold^18,19^. These structures provide a foundational understanding of the C1 and C2 microtubules. However, the dissociation of C1 and C2 microtubules during the separation and purification process of the CA sample leads to the disruption the intact CA density map. Consequently, essential structural details are absent, particularly for components situated at the bridge region and those interacting with both C1 and C2 microtubules. This results in a limited understanding of the C1 and C2 connections in the asymmetrical assembly of CA. Moreover, the assembly of CA in higher animals remains unknown due to significant differences in CA component proteins between lower species and higher animals^20^.

In this study, we employed the cutting-edge visual proteomics technology^21^, combining cellular cryo-electron tomography (cryo-ET) and AlphaFold2 modeling, determined the near-complete native structure of the intact CA from mouse sperm. By employing subtomogram averaging (STA), reconstruction of the CA structure was achieved with a resolution of up to 5.5 Å. The well-resolved tertiary structures enabled us to build models of 39 different CA-associated proteins using AlphaFold2 predictions, including both homologous and newly identified component proteins. Notably, our study revealed the full-length structures of the long chain-like proteins CFAP47 and HYDIN, providing insights into their molecular mechanisms in connecting C1 and C2 microtubules. Additionally, we found that sperm from *Cfap47*-knockout (KO) mice exhibited a hollowing of the CA central bridge, correlating with their reduced progressive motility. It provides an example to investigate the molecular pathogenesis of CA related mutations directly from individual samples by cryo-ET approach. In conclusion, the visual proteomics approach^21^ enabled the identification of novel CA components in mammalian sperm without the need for cell disruption or biochemical purification. The structural insights gained are essential for understanding the molecular mechanisms of CA components in ciliary motility, and the implications of their mutations in human ciliopathies.

## Results

### Overall structures of mouse sperm CA

The freshly extracted mouse sperm were vitrified on EM grids. To facilitate cryo-ET imaging, lamellae with approximately 200 nm thickness were generated by cryo-FIB milling (Fig. S1). Tilt series were recorded using a 300 kV Krios cryo transmission electron microscope (TEM) and a dose-symmetric scheme with a tilting increment of 3 degrees. The structural features of the CA particles are clearly visible in the tomograms, and their positions and orientations (two of three Euler angles) can be confidently determined (Fig. S2). By employing solely data-driven templates, three-dimensional (3D) classification and refinement of subtomograms revealed the 32 nm-repeating units of CA (Fig. S2). When we enlarged the box size of the CA map to cover the entire “9+2” axoneme complex, we could observe the other components such as DMT, ODA, IDA, N-DRC, and RS (Fig. 1A). It indicates that the structural integrity of the entire axoneme has been preserved in our sample, and the positioning of CA within the axoneme remains stable.

**Figure 1.**
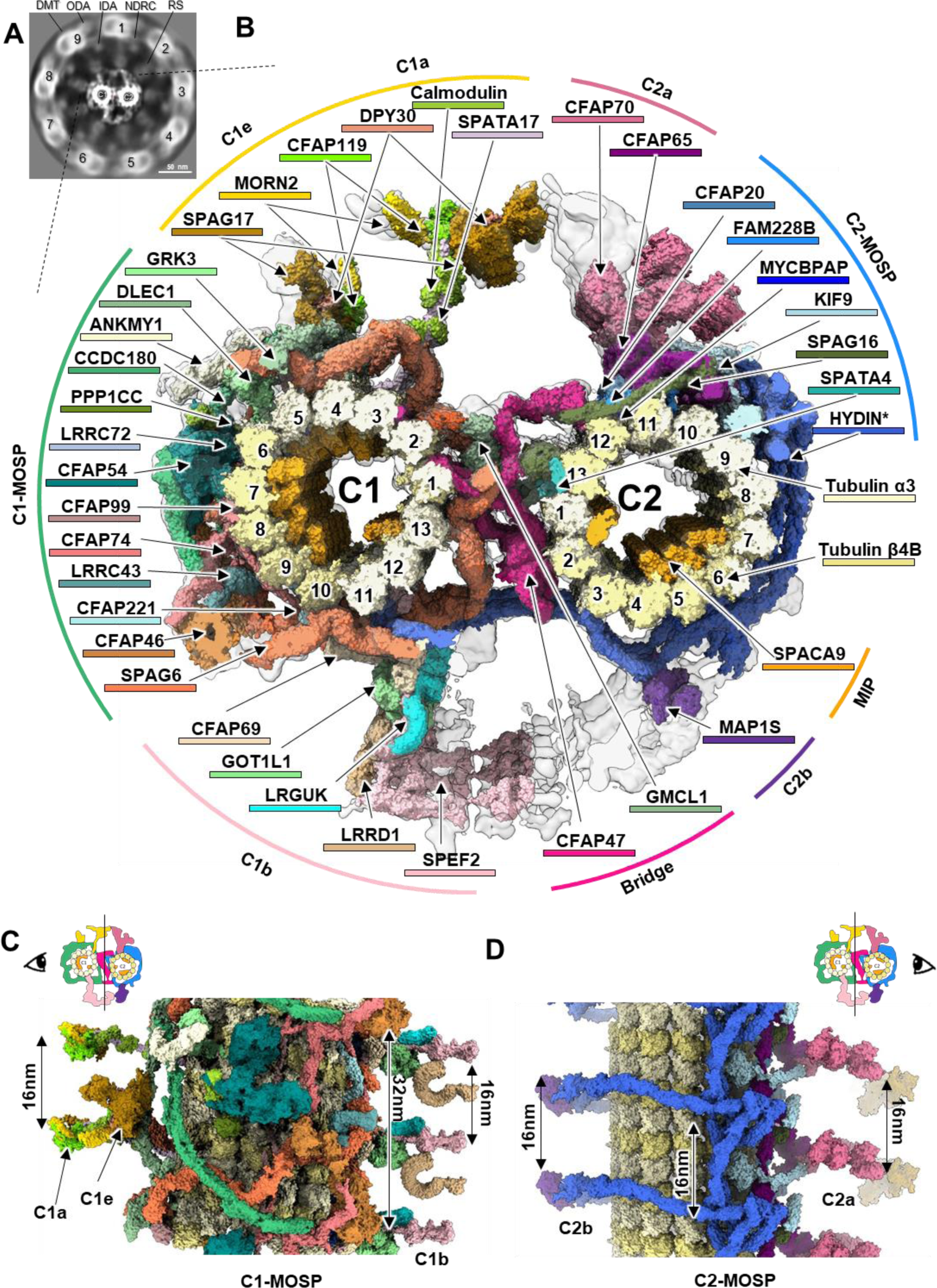
Overall structure of mouse sperm CA. (**A**) Slice view of the entire mouse sperm axoneme structure, highlighting the major components: DMT, RS, ODA, IDA, N-DRC. The numbers indicate the components in different directions. (**B**) Molecular composition of the CA architecture. The protein components are categorized based on their specific locations within the CA, and are colored differently (Table S4). (**C**) The side view of C1 half of the CA, displaying both 16 nm and 32 nm repeats. (**D**) The side view of C2 half of the CA, displaying predominant 16 nm repeat.

To address the structural flexibility for this huge complex, local refinements were performed on distinct regions of the CA map, resulting in a resolution of 7.7 Å for the 32 nm repeats of C1 and 6.6 Å for the 16 nm repeats of C2. In addition, the 16 nm repeats of bridges achieved a resolution of 7.7 Å, 8 nm repeats of C2 achieved 5.5 Å resolution, and projections exhibited resolution ranges from 7.8 Å to 18 Å (Fig. S2 & S3). Individual α-helices for the tubulins and multiple associated proteins were resolved in these maps. By combining local maps, we generated a composite reconstruction of the native CA structure.

This composite map was subsequently compared to published cryo-EM maps of CA from various species (Fig. S4 & S5). Notably, the cryo-EM map of isolated CA from *C. reinhardtii* cilia, obtained through single-particle cryo-EM^18,19^, exhibited a severe loss of connection densities between C1 and C2 microtubules, including the bridge, C1a/C2a, and C1b/C2b regions (Fig. S4A). Additionally, there were density losses around C2 and C2b projections, whereas our map preserved these densities entirely (Fig. S4). We also noted a significant shift in the projection of C1b and C2a in their maps compared to ours, likely due to the absence of a pulling force from C2b and C1a in the isolated sample (Fig. S4). Furthermore, the C1f projection in lower species disappear in higher species, consistent with previous reports^22^ (Fig. S4A). The resolutions of the in situ structural studies of CA reported so far are all relatively low, below 26 Å ^22,23^(Fig. S5). Our density map aligns closely with these maps, confirming the accuracy of our structural analysis as well as the conservation of overall CA structure across different species. Notably, we observed that the overall structures of human sperm and mouse sperm are highly similar, with the only difference being the absence of some microtubule inner proteins (MIPs) in the C2 lumen of human sperm (Fig. S5A).

The clearly resolved secondary and tertiary structures in most regions of our CA map enabled us to interpret maps and build pseudo-atomic models (Fig. S6-S18). This was accomplished by incorporating various visual proteomics methods^21^, including using homologue structural data from *C. reinhardtii* cilia CA^18,19^, conducting mass spectrometry (MS) of mouse sperm (Table S2)^24^, screening AlphaFold2-predicted models with DomainFit tool^25^, performing mutagenesis studies (refer to Methods for more details). Our final model of 32 nm repeat in the CA structure of mouse sperm contains 260 tubulins and 206 non-tubulin chains, which correspond to 39 identified proteins and 6 unknown proteins (Table S2 & S3).

The overall structure of mouse sperm CA is an asymmetric complex with distinct projections around the C1 and C2 microtubules (Fig. 1B & Video S1). Both microtubules consist of 13 protofilaments with seams that face each other at the connecting bridge between them. All CA sub-complexes are interconnected through intricate networks of scaffolding and connecting proteins. In addition to the C1 and C2 microtubules, we categorized the organization of mouse sperm CA into four groups: (1) a network of tightly bound microtubule outer surface proteins (MOSPs) in C1 and C2, (2) five large projections (C1a/C1e, C1b, C2a, C2b), (3) MIPs in C1 and C2 lumens, and (4) bridge that connects C1 and C2 (Fig. 1B). C1a, C1b, C2a and C2b projections in CA all exhibit 16 nm repeats, while the C1e projection exhibit a 32 nm repeat (Fig. 1C & 1D). The C1-MOSP majorly shows 32 nm repeats, while the C2-MOSP and bridge predominantly exhibits 16 nm repeats (Fig. 1C & 1D).

### Assembly of C1-MOSP

The rapid beating behavior of cilia requires the CA to have structural elasticity and stability. The primary component of C1-MOSP is sperm-associated antigen 6 (SPAG6), which forms a spring-shaped scaffold around the C1 microtubule (Fig. 2A). SPAG6, a highly conserved armadillo-repeat protein homologous to PF16 in *C. reinhardtii*, serves as a fundamental building block, polymerizing on the outer surface of C1 to form stable spirals in both 32 nm and 16 nm repeats (Fig. 2A). The structure of the SPAG6 monomer and the spring scaffold polymer is identical to the reported structure of the CA in *C. reinhardtii* cilia^18^, indicating that this structure is highly conserved and crucial for the CA’s assembly and function. This spring scaffold potentially acts like a mechanical spring, providing the CA with the necessary structural elasticity and stability essential for ciliary beating. Deficiency in SPAG6 has been associated with male infertility (impaired sperm motility) and hydrocephalus^26^ (Fig. S19A & Table S5).

**Figure 2.**
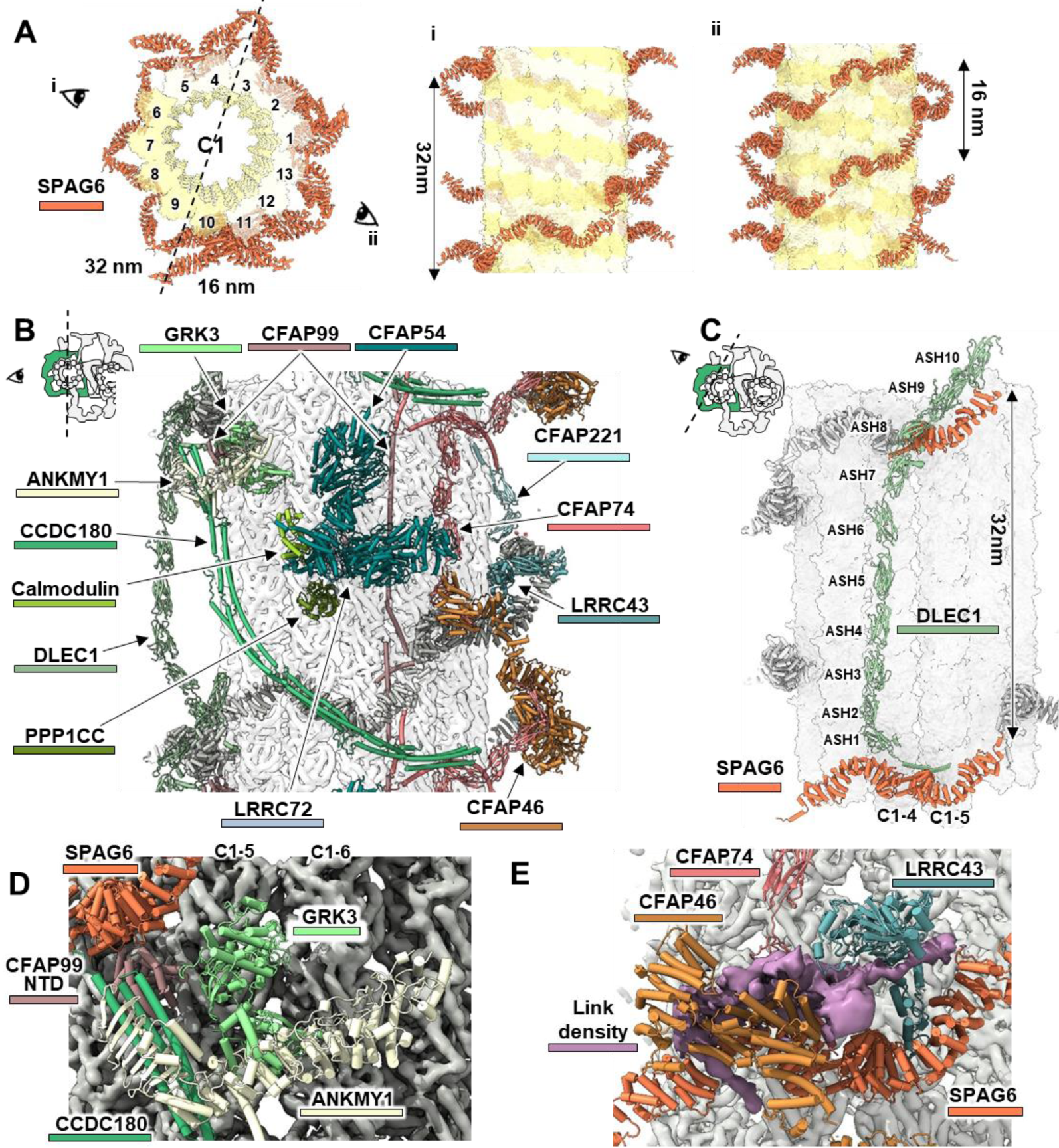
Structural composition of C1-MOSP in mouse sperm CA. (**A**) SPAG6 is identified as the primary component of C1-MOSP, forming 16 nm and 32 nm repeats around the C1 microtubule. This is displayed in one cross-sectional view and two side views. (**B**) Depiction of the overall protein composition of C1-MOSP in the 32 nm repeat. Proteins are distinctly colored for clarity (Table S4). (**C**) The ASH protein DLEC1 interacts with two SPAG6 proteins via its NTD and CTD, establishing the 32 nm periodicity. (**D**) Detailed view within C1-MOSP highlighting the assembly of CFAP99, GRK3, ANKMY1, and CCDC180. (**E**) Detailed view within C1-MOSP highlighting the assembly of CFAP46, CFAP77, LRRC43, and SPAG6, along with an undefined link density (pink).

On the outer wall of C1, SPAG6 exhibits a 32 nm periodicity, matching the distance between adjacent RSs in the DMT. Within this 32 nm interval, more than a dozen distinct components of CA are arranged in a well-ordered manner (Fig. 2B). Proteins containing the ASH domain (<u>A</u>SPM, <u>S</u>PD-2, <u>H</u>YDIN) are commonly associated with the CA asembly^27^. The protein Deleted in Lung and Esophageal Cancer 1 (DLEC1) has 10 ASH domains and interacts with SPAG6 at 32 nm intervals through its N-terminal domain (NTD) and C-terminal domain (CTD), contributing to this periodic structure (Fig. 2C). The N-terminal helix and ASH1 domain of DLEC1 interact with the adjacent SPAG6 dimer, while its C-terminal ASH8-10 domains bind to a single SPAG6 in the next 32 nm repeat (Fig. 2C). The central region of the DLEC1 protein is fully extended and aligned almost parallel to protofilament-4 of C1 tubule, forming the structural foundation of the C1e projection. Mutations in FAP81, the homologous protein of DLEC1 in *C. reinhardtii* cilia, result in the absence of C1e projection ^28^.

In addition to SPAG6, several other proteins are distributed along the longitudinal axis of the microtubule on the outer wall of C1. Specifically, CCDC180 and CFAP99 are predominantly composed of long α-helices, while CFAP221 contains ASH domains. CFAP74 features both long helices and multiple ASH domains. These proteins not only contribute to the formation of the 32 nm periodic structure but also serve as scaffolds for the recruitment of other globular proteins, such as ANKMY1(Ankyrin repeat and MYND domain-containing protein 1), PPP1CC (Serine/threonine-protein phosphatase PP1-gamma catalytic subunit), LRRC43(Leucine-rich repeat-containing protein 43), GRK3(G protein-coupled receptor kinase 3), CFAP54, LRRC72, and CFAP46 (Fig. 2B). Notably, CFAP54 and CFAP46, the two largest globular proteins, exhibit distinct structural differences compared to their homologs in *C. reinhardtii* cilia (Fig. S20A&B). The NTD of CCDC180 interacts with ANKMY1, GRK3, and the N-terminal globular domains of CFAP99 (Fig. 2D). CFAP54 encases a calmodulin and subsequently binds to LRRC72 and PPP1CC (Fig. S20C). Additionally, CFAP46 attaches to SPAG6, LRRC43, and CFAP74 through an unspecified linkage (Fig. 2E). These proteins are situated close to the heads of RS8 and RS9, indicating their potential involvement in CA-RS interactions (Fig. 1A). Several mutations in these proteins have been linked to human PCDs (Table S5), such as CFAP54 and CFAP74 (Fig. S19B & C), proving their important roles in maintaining CA function.

In C1-MOSP, several CA components specific to mammals, such as ANKMY1, GRK3, and LRRC43, differ from the CA structure observed in *C. reinhardtii*^18,19^. ANKMY1, GRK3 are positioned near the RS8 interface with CFAP54, while LRRC43 seems to interact with RS7 with CFAP46 (Fig. 1A & B). Beyond contributing to the CA-RS interface, these components may have additional biological functions. Notably, GRK3 exhibits phosphorylation activity; although its presence as a CA component is somewhat unexpected (Fig. S21). Phosphorylation is known to provide critical chemical signals regulating ciliary motility^12^, suggesting that PPP1CC and GRK3 within the C1-MOSP may be involved in this regulatory process.

### Assembly of large C1-projections

C1 has three large projections: C1a, C1b, and C1e. Although the local resolutions in these regions are not high enough for direct model building, their overall structures resemble those of *C. reinhardtii* CA^18,19^. Consequently, homologous structure information and AlphaFold2 Multimer predictions can assist in protein identification and model building (Fig. S6). C1a and C1e are closely positioned and share common components, including SPAG17, MORN2, CFAP119, and DPY30. Both C1a and C1b feature narrow microtubule-bound stalks and large distal heads (Fig. 3).

**Figure 3.**
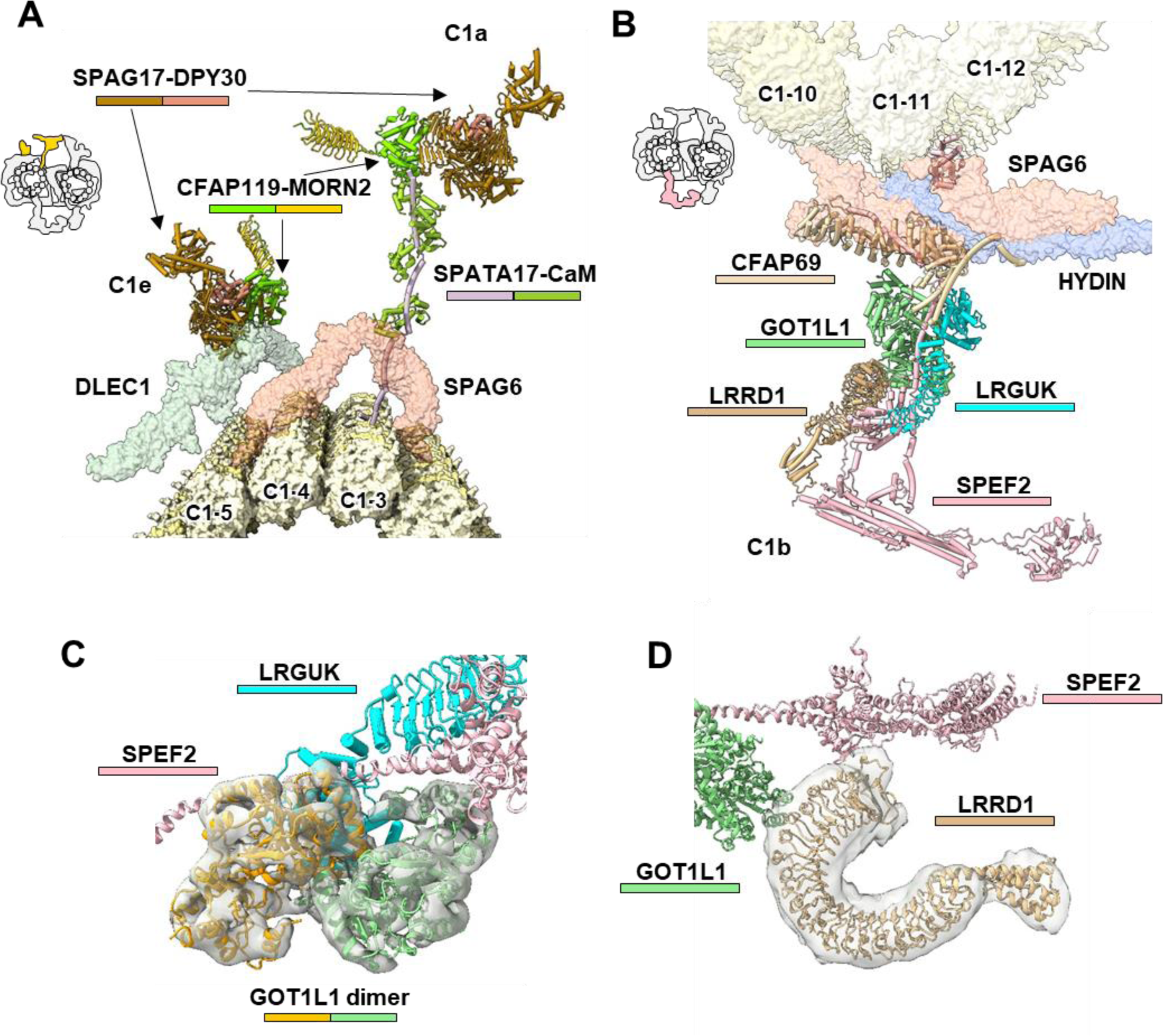
Structural composition of C1 projections in mouse sperm CA. (**A**) Protein composition of the C1e and C1a projections. (**B**) Protein composition of the C1b projection. (**C**) Location and structure of the newly identified GOT1L1 dimer in the C1b projection. (**D**) Location and structure of the newly identified LRRD1 in the C1b projection. Proteins are distinctly colored for clarity (Table S4).

The assembly of the C1a projection centers around the rachis protein SPATA17, which extends from the C1 protofilament-3 and binds to SPAG6 (Fig. 3A). Three calmodulin densities are located on the SPATA17 helical region, forming the stalks for the C1a projection (Fig. 3A), indicating the importance of calcium in regulating the conformation or dynamics of the C1a projection. The SPAG17-DPY30 complex constitutes the head of both C1a and C1e projections (Fig. 3A).

Compared to its homolog PF6 in *C. reinhardtii*, SPAG17 possesses an additional globular domain and several longer helices (Fig. S21A). In *Spag17*-KO mice, the C1a projection disappears in the axoneme CA^29^, consistent with our structural observations. FAP114 and FAP119 in *C. reinhardtii* share a homologous protein CFAP119 in mouse (Fig. S22). Besides, the helical bundles formed by the algal-specific protein FAP7, the α-helical connection networks between C1a and C1e, and the stalk-attached protein FAP101 in *C. reinhardtii*^18,19^ are all absent in our density map (Fig. S21B), highlighting evolutionary differences in C1a projections.

The assembly of the C1b projection centers around the rachis protein SPEF2 (CPC1 in *C. reinhardtii*), which protrudes from the C1 protofilament-11/12 and interact with SPAG6 and HYDIN NTD (Fig. 3B). The recruitment and positioning of SPEF2 are critical for the formation of the C1b projection, while mutations in *Spef2* gene lead to immotile sperm and PCD^30^. The extended SPEF2 organizes a large cluster of C1b projection proteins including CFAP69, GOT1L1 (putative aspartate aminotransferase), LRRD1 (leucine-rich repeat and death domain-containing protein 1), and LRGUK (Fig. 3B-D). Notably, SPEF2 and LRGUK exhibit structural differences compared to their homologs in *C. reinhardtii* cilia (Fig. S23A & B). SPEF2 and LRRD1 are located near the RS6 interface (Fig. 1A) and seem to participate in CA-RS interaction. Several mutations in C1b projections have been linked to human PCDs (Table S5), such as CFAP69 and SPEF2 (Fig. S19D & E), proving their important roles in maintaining CA function.

Proteins of FAP42, HSP70, and enolase, which have been reported in the C1b structure of *C. reinhardtii* CA^18,19^, are not present in our map (Fig. S23C). Aside from the guanylate kinase activity of LRGUK, their enzymatic and chaperone activities appear to be lacking in the C1b projections in mammals. These functions could be performed by unidentified proteins in mammalian CA or might have been lost during evolution. The identification of GOT1L1 in sperm CA is particularly noteworthy (Fig. 3C). GOT1L1 is highly expressed in mouse testis and is potentially involved in D-aspartate synthesis^31^. The levels of D-aspartate in mouse testis are regulated spatiotemporally and play a role in modulating spermatogenesis^32^. The localization of GOT1L1 in sperm CA supports its stable and ongoing function in D-aspartate synthesis.

### Assembly of C2-MOSP and C2 projections

C2-MOSP exhibits distinctive features compared to C1-MOSP. The C2 microtubule lacks PF16 spirals and shows periodicities of both 16 nm and 8 nm. The 16 nm periodicity predominantly includes SPAG16, KIF9 and HYDIN (Fig. 4A). SPAG16 features a C-terminal β-propeller domain that binds with an 8 nm periodicity to the seam of the C2 microtubule and a coiled-coil domain that dimerizes with adjacent copies, resulting in a transition from 8 nm to 16 nm periodicity (Fig. 4A). The SPAG16 dimeric helix forms the structural foundation of CFAP65 and the C2a projection. The β-propeller domain of SPAG16 binds with SPATA4 at the seam (Fig. 4A). SPAG16 is crucial for CA assembly, as the entire CA is lost when its homolog protein PF20 is mutated in *C. reinhardtii*^33^. CFAP20 are well-known to connect the A and B tubules in the DMT^34^, but in CA it attaches to the C2 microtubule through a distinct mechanism with minimal contact with the tubulin surface (Fig. 4A). Compared to the *C. reinhardtii* CA structure, SPATA4 has an additional CTD compared to its homolog FAP178 (Fig. S24A), which seems to enhance its interaction with the C2 microtubule wall. Similarly, FAM228B has an additional CTD compared to its homolog FAP239 in the *C. reinhardtii* CA (Fig. S24B), enhancing its interaction with the C2 microtubule.

**Figure 4.**
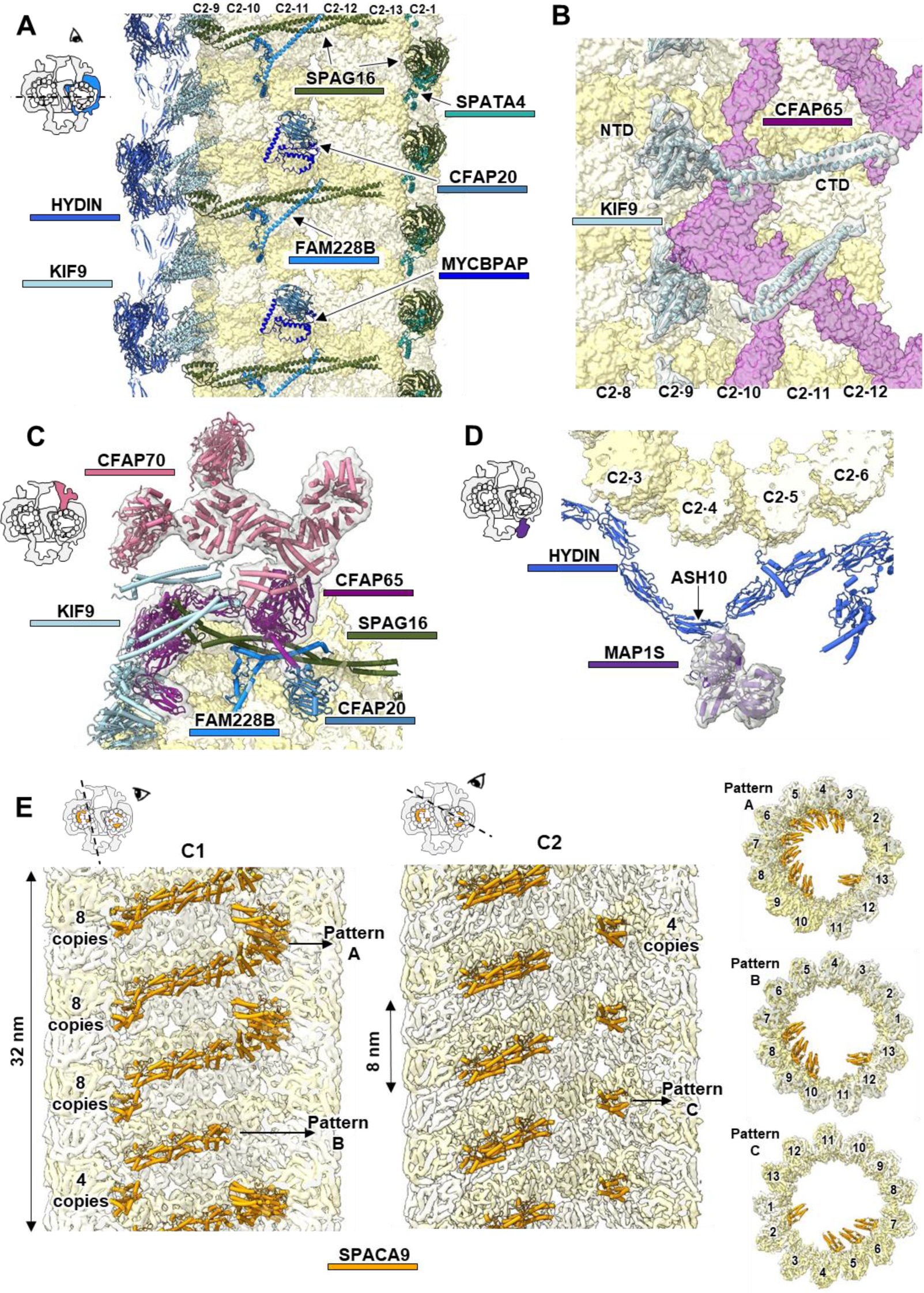
Structural composition of C2 microtubule and MIPs in mouse sperm CA. (**A**) Depiction of the overall protein composition of C2-MOSP in the 16 nm repeat. (**B**) Location and structure of the KIF9 NTD and CTD in the C2-MOSP. (**C**) Location and composition of the C2a projection. (**D**) Location and composition of the C2b projection. (**E**) Structural composition of MIPs in mouse sperm CA. SPACA9 exhibits 32 nm and 8 nm repeats in the lumen of the C1 and C2 microtubules, respectively. SPACA9 showcases three distinct patterns on the microtubule wall within the CA. Proteins are distinctly colored for clarity (Table S4).

Similar to SPAG16, KIF9 has an N-terminal globular domain that binds with 8 nm periodicity to protofilament 9 of the C2 microtubule, and a C-terminal coiled-coil domain that binds to different regions of CFAP65 in various conformations, leading to a shift from 8 nm to 16 nm periodicity (Fig. 4B). Mutations in *KIF9* significantly impair sperm motility^35^, underscoring its critical role in CA function. Notably, the CTD of KIF9 was not identified in prior structures of *C. reinhardtii* CA^18,19^, whereas our in-situ structure provides more complete structural information.

We identified two major components in C2a projections: CFAP65 and CFAP70. CFAP65 is situated on a structural base consisting of KIF9, FAM228B, CFAP20, and SPAG16 (Fig. 4C). CFAP70 forms a homodimer and is located on top of CFAP65. CFAP70 is positioned near the RS2 interface (Fig. 1A) and appears to participate in the CA-RS interaction. Several mutations in C2-MOSP and C2a projection have been linked to human PCDs (Table S5), such as CFAP20, SPAG16, KIF9, CFAP65 and CFAP70 (Fig. S19F & G), proving their important roles in maintaining CA function.

The C2b projection is the least understood part of the CA structure. Due to its highly dynamic nature, its structural composition and functions remain speculative. Here, we identified HYDIN as the main component of this region (discussed later), and also found that MAP1S (microtubule-associated protein 1S) is located in C2b (Fig. 4D). According to the Human Protein Atlas database, MAP1S is highly expressed in spermatids, but its exact function is not yet known. Our structure shows that MAP1S binds to the ASH10 domain of the HYDIN protein, forming the core structure of the C2b projection, potentially facilitating the assembly of other C2b components (Fig. 4D). Further studies are required to obtain more complete structural information on the C2b projection to elucidate the detailed function of MAP1S.

### The primary component of MIP in mammalian CA is SPACA9

Unlike the abundant MIPs found in the DMT A and B tubules^34^, the C1 and C2 microtubules in the CA exhibit lower MIP density. The most prominent MIP in the C1 and C2 microtubules is the regularly arranged protein SPACA9 (Fig. 4E). SPACA9 has a simple structure composed of eight α-helices organized into a four-helix bundle, with four shorter helices inserted into it. These shorter helices cap the bundle and form the microtubule-binding interface. Recent studies on human sperm microtubule singlets (SMT) and DMT have shown that SPACA9, with its pyramid-shaped conformation, decorates the lumens of DMT B-tubule and SMT with specific repeating patterns^34,36^. For instance, in the DMT of human tracheal cilia, SPACA9 exhibits an 8 nm periodicity and is situated between α- and β-tubulin heterodimers^36^. In contrast, in the lumens of the C1 and C2 microtubules in mouse sperm CA, SPACA9 displays two new arrangements with 48 nm and 8 nm periodicities, respectively (Fig. 4E).

In the C1 lumen, SPACA9 forms an 8-8-8-4 pattern within each 32 nm repeat (Fig. 4E). For the eight SPACA9 copies (pattern A) per layer, seven copies are continuously located between protofilaments 3 and 10, while one isolated SPACA9 is found between protofilaments 12 and 13. For the four SPACA9 copies (pattern B) per layer, three copies are consistently between protofilaments 7 and 10, with one isolated SPACA9 between protofilaments 12 and 13. In the C2 lumen, SPACA9 forms an 8 nm repeating pattern (pattern C) with four copies per layer. In this pattern, one isolated SPACA9 is between protofilaments 1 and 12, and the remaining three copies are between protofilaments 4 and 7. The positioning of SPACA9 at the intersection of tubulin heterodimers and various protofilaments contributes to the stabilization of the tubule walls. Pattern B and pattern C display a mirror symmetry centered around the bridge, reinforcing the binding sites for RS7 and RS4, respectively (Fig. 1A). Additionally, pattern A enhances the binding sites for RS8 and RS9 (Fig. 1A).

The MIPs in the C1 and C2 microtubules of mammalian CA differ considerably from those in *C. reinhardtii* CA^19^, where the MIPs predominantly consist of FAP196, FAP225, and several nonglobular MIPs (Fig. S25). In *C. reinhardtii* CA’s C1 microtubule, FAP275 and FAP105 are the major MIPs, which do not overlap with SPACA9 in mouse CA (Fig. S25A). In the C2 microtubule of *C. reinhardtii* CA, FAP196 occupies a position that overlaps with SPACA9 in mouse CA (Fig. S25B). The homolog of FAP196 in mice, CFAP52, functions in microtubule stabilization in DMT^34^, suggesting that SPACA9 replaces CFAP52 in higher animals’ C2 microtubules for stabilization. Other MIPs in *C. reinhardtii* CA’s C2 microtubule, such as FAP239, FAP388, FAP225, FAP213, and FAP424, do not conflict with SPACA9 in mouse CA (Fig. S25B). These findings suggest that in higher animals, the roles of these MIPs might be integrated and simplified, with SPACA9 predominantly enhancing microtubule stability. Furthermore, there are also some flexible MIPs in the C1 and C2 lumens of mammalian CA, mostly distributed at the tubulin binding interface. Due to the absence of distinct globular domains, it is impossible to determine their sequences and identify the respective proteins in our map.

### CFAP47 and GMCL1 form the CA bridge

In CA structure, C1 and C2 are connected at three principal regions: a central bridge that links the two microtubules and two lateral connections between the opposite projections of C1a/C1b and C2a/C2b at the periphery (Fig. 1B). While CFAP47 and HYDIN, two ASH domain-containing proteins, have traditionally been implicated in the C1-C2 linkages, their exact roles have remained ambiguous due to the limited structural information available for these proteins. Based on our near-complete in situ model of mammalian CA, which includes the full-length structures of CFAP47 and HYDIN, we discovered that these proteins are essential and function in a complementary manner in tethering the C1 and C2 microtubules.

CFAP47 forms the structural backbone of the CA bridge (Fig. 5A). It possesses 20 ASH domains (Fig. S26), with its NTD attaching to the C1 microtubule and extending to the C2 microtubule via its middle domain and CTD (Fig. 5B). The ASH1 and ASH4 domains of CFAP47 bind to protofilament-13 and protofilament-1 of the C1 microtubule, respectively, reinforcing the C1 seam. From ASH5, CFAP47 bends back to stretch towards the C2 microtubule. The ASH6-10 domains, along with a calponin homology (CH) domain consisting of α-helices, form a large and stable structure called the CFAP47-ring (Fig. 5B). This CFAP47-ring binds to protofilament-2 of the C2 microtubule through its ASH7 domain and interacts with the ASH5-6 domains of HYDIN via its ASH8-9 domains (Fig. 5C & S27). Subsequently, the remaining CFAP47 domains extend laterally along the surface of the C2 microtubule, with the ASH13 domain forming stable interactions with its ASH3 domains, thereby reinforcing the connections between CFAP47’s NTD and CTD, and enhancing the linkage between C1 and C2 microtubules (Fig. 5D & S28). The ASH18 domain of CFAP47 binds to SPAG16, and its C-terminal ASH20 domain binds to protofilament-12 of the C2 microtubule (Fig. 5E). In conclusion, the extensive interaction network established by CFAP47, including internal interactions among its ASH domains and external connections with the C1/C2 protofilament at the seams, highlighting its critical roles in CA assembly.

**Figure 5.**
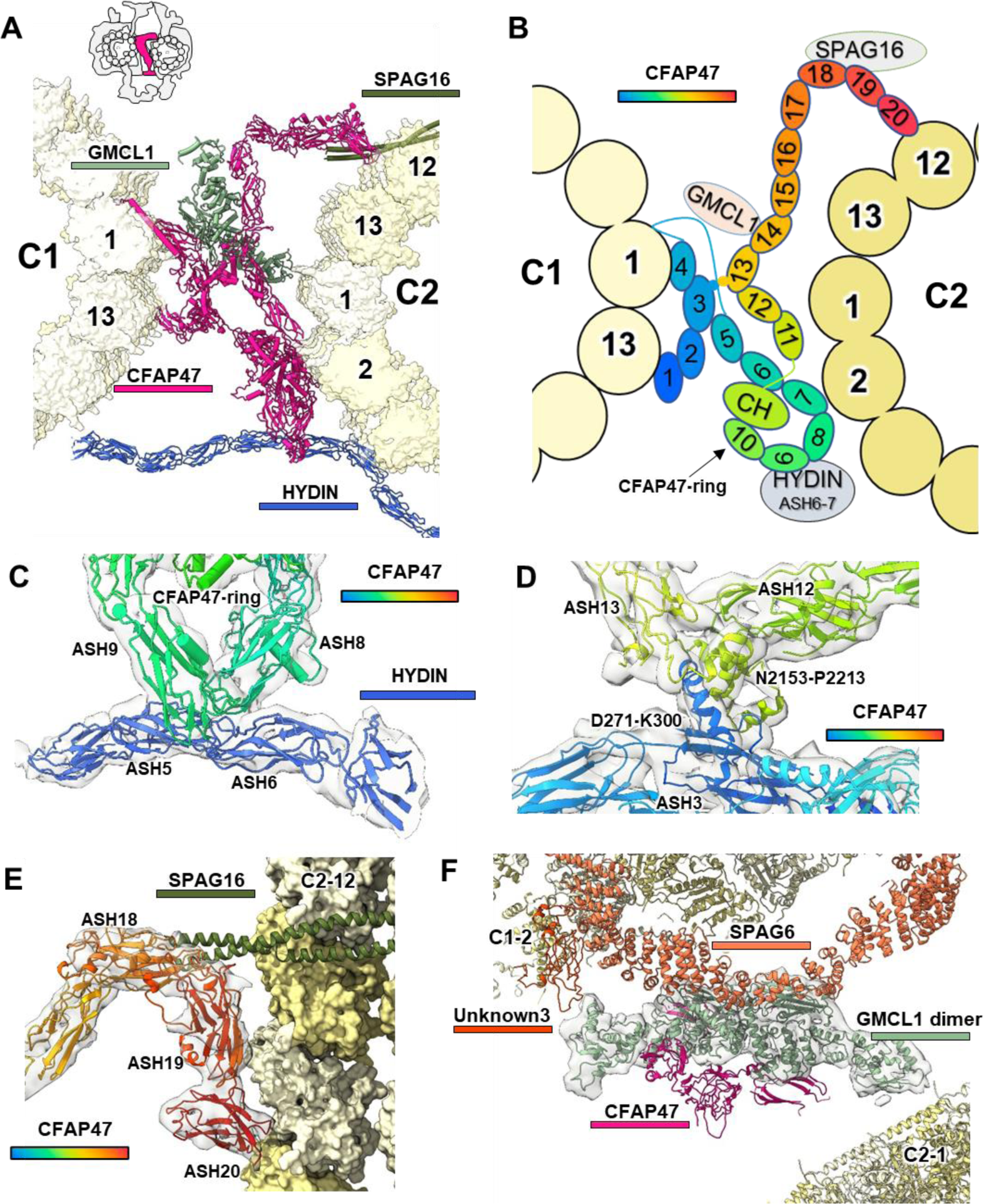
Structural composition of the bridge in mouse sperm CA. (**A**) Depiction of the overall protein composition in CA bridge within 16 nm repeat. (**B**) Schematic representation of all CFAP47 domains and their interactions in the bridge. The number in CFAP47 structure indicates the sequence position of the ASH domains. (**C**) Detailed view of the interactions between CFAP47 and HYDIN. (**D**) Detailed view of the interactions between the ASH3 and ASH13 domains of CFAP47. (**E**) Detailed view of the interactions between the CFAP47 CTD and the C2 microtubule. (**F**) Localization and structure of the newly identified GMCL1 dimer within the bridge.

In addition to CFAP47, we identified the germ cell-less 1 (GMCL1) protein situated within the CA bridge and closely associated with CFAP47 (Fig. 5A). GMCL1 is a protein related to spermatogenesis and asthenozoospermia (AZS)^37^, yet its function remains undefined. Our findings reveal that GMCL1 is a globular protein primarily composed of α-helices, with its CTD forming homodimers (Fig. 5F). GMCL1 facilitates interactions between SPAG6 in C1-MOSP and ASH14 of CFAP47, thereby strengthening the binding of CFAP47 to the C1 microtubule (Fig. 5F). Additionally, the NTD of GMCL1 interacts with an unidentified protein (termed Unknown3) located on protofilament-2 of the C1 microtubule (Fig. 5F), indicating its role in enhancing the structural integrity between C1 and C2 microtubules.

### HYDIN externally connects C1 and C2

Unlike CFAP47, which connects the seam areas of C1 and C2 in a highly folded chain structure in the middle of CA, the 33 ASH domains of HYDIN (Fig. S26) form a much-extended chain structure encircling C1 and C2 from the outside in a semicircular manner (Fig. 6A). HYDIN and CFAP47 directly interact with each other, effectively connects C1 and C2 from both external and internal positions (Fig. 6A). Interestingly, the HYDIN protein utilizes its ultra-long chain structure to reinforce the CA architecture in both radial and axial directions. The radial reinforcement is mediated by the ASH1-18 domains, while the axial reinforcement involves the ASH19-33 domains (Fig. 6A & B). The N-terminal ASH1 domain of HYDIN starts from the protofilament-11 of the C1 microtubule, surrounded by SPAG6, CFAP69, and SPEF2, and proceeds towards the C2 microtubule (Fig. 6C). In the bridge, the ASH6-7 domains of HYDIN interact with the ASH8-9 domains of the CFAP47-ring, leading to the binding of the two longest ASH proteins in CA (Fig. 5C). The ASH8 domain of HYDIN then attaches to the protofilament-3 of the C2 microtubule, curving its ASH8-12 domains to form the scaffold of the C2b projections. The ASH13-18 domains subsequently extend laterally along the C2 microtubule to the protofilament-8 (Fig. 6A).

**Figure 6.**
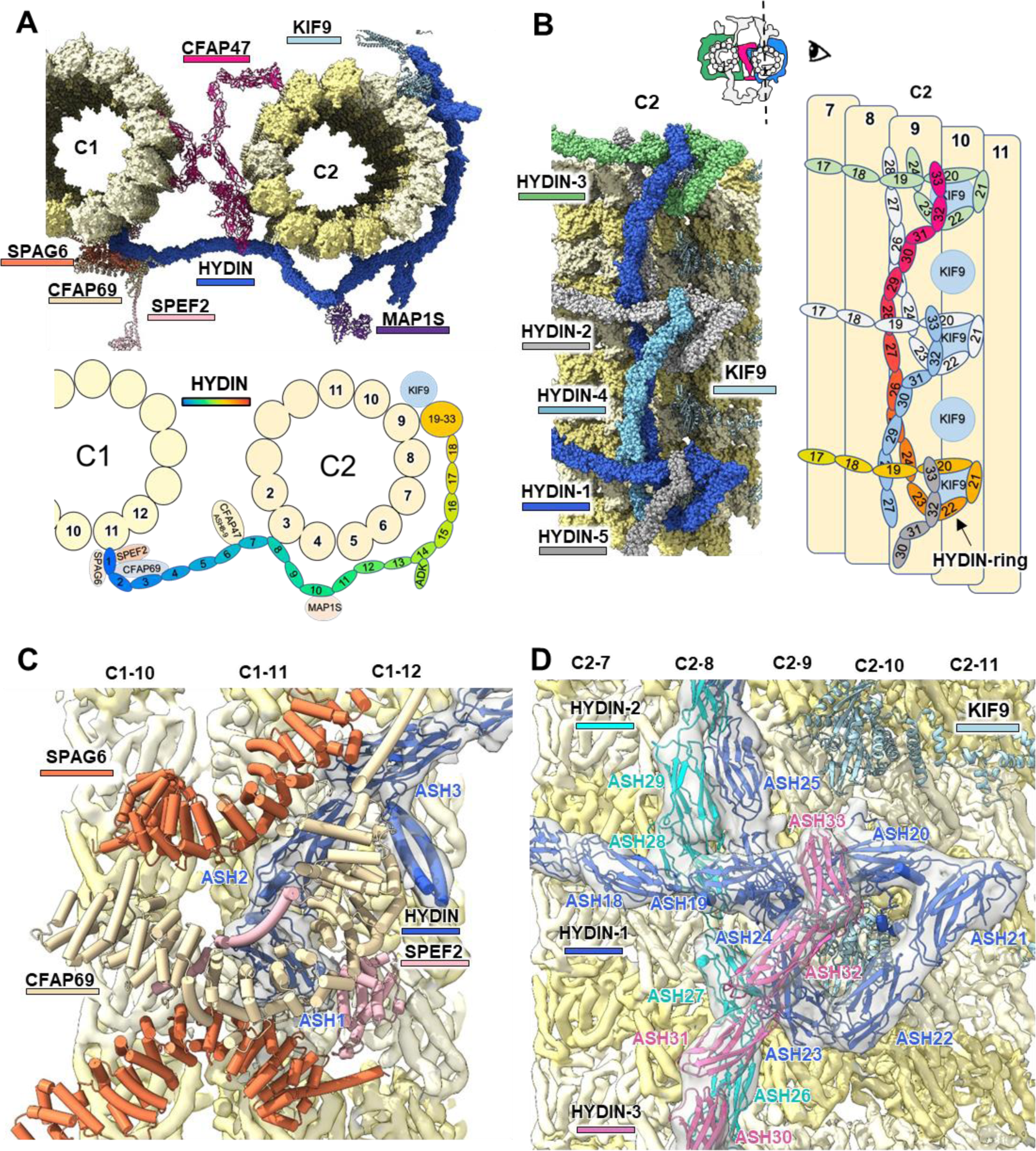
Structure of HYDIN in mouse sperm CA. (**A**) Overall structure and schematic representation of HYDIN, shown in cross-sectional view. Numbers within the HYDIN structure indicate the sequence position of the ASH domains. (**B**) Structure and schematic representation of the HYDIN CTD in side view, highlighting ASH17-33 domains. Different HYDIN proteins are shown in distinct colors. (**C**) Detailed view of the interactions between the HYDIN NTD and the C1 microtubule. (**D**) Detailed view of the interactions within the HYDIN CTD trimer on the C2 microtubule. Three HYDIN proteins are displayed in different colors, with their ASH domains labeled.

Next, the ASH19-33 domains of HYDIN form a unique 270-degree bend and extend longitudinally along with C2 protofilaments (Fig. 6B). Notably, the remaining HYDIN domains are over 32 nm in length, enabling three HYDIN proteins to overlap within every 16 nm interval, thereby creating a HYDIN trimer complex (Fig. 6B & S29). Specifically, the ASH19-24 domains of HYDIN-1 form a triangular ring known as the HYDIN ring, with the ASH25-28 domains passing through the gap between the HYDIN-2 ring and the C2 microtubule, and the ASH29-33 domains settle on top of the HYDIN-3 ring (Fig. 6B). Consequently, each HYDIN ring is clamped by two other HYDIN molecules from 16 nm and 32 nm intervals away, forming a highly compact complex known as the HYDIN-ring complex (Fig. 6D). This HYDIN-ring complex binds to the KIF9 globular domain on the C2 microtubule (Fig. 6D). The presence of HYDIN-ring complex in other low-resolution CA models suggests that this architecture is highly conserved across species (Fig. S4 & S5).

The full-length structure of HYDIN we present is consistent with previous functional evidence. Biochemical analyses demonstrate that HYDIN interacts with CPC1 (SPEF2 in mouse) and kinesin-like protein 1 (KLP1, KIF9 in mouse) ^38^, while these direct interactions are both observed in our structure. It has also been reported that in spermatozoa lacking HYDIN, KIF9 is undetectable^35^, which may be related to the interactions between HYDIN and KIF9 on the CA. RNA interference knockdown of HYDIN results in short flagella lacking the C2b projection^38^, underscoring the essential role of HYDIN as a structural component of C2b. Furthermore, we identified the adenylate kinase-like domain (ADK) domain of HYDIN in the C2b projections (Fig. 6A), which catalyzes the interconversion of various adenosine phosphates. This ADK domain is located near the RS4 head interface (Fig. 1A) and may be involved in the CA-RS regulatory role for nucleotides, given the discovery of nucleotide-binding domains in the RS heads^39,40^. As axonemes consume approximately 230,000 molecules of ATP per beat cycle^41^, they may be sensitive to local fluctuations in the ATP/ADP ratio and regulated by the adjacent HYDIN-ADK domain and RS4 head. Many mutations in *Hydin* gene have been linked to human PCDs (Fig. S19H & Table S5), proving its important roles in maintaining CA structure and function.

### *Cfap47*-KO sperm exhibit reduced progressive motility and impaired bridge density

The deficiency of HYDIN has been reported to cause congenital hydrocephalus in mice and lead to fetal death^42^, highlighting the critical role of the HYDIN protein in CA assembly. This observation aligns with our structural findings, which show that the HYDIN protein is involved in the assembly of C1b, bridge, C2b, and C2-MOSP (Fig. 6). However, the impact of CFAP47 deficiency on CA architecture and its mechanism to cause abnormal sperm motility remains ambiguous. To address this, we generated a *Cfap47*-KO mouse strain by targeting exons 2‒50 of the *Cfap47*-205 (ENSMUST00000197180.6) transcripts using CRISPR-Cas9 technology (Fig. S30A). Sanger sequencing confirmed the successful deletion of 171,604 base pairs (7,249 bp coding sequence) (Fig. S30B). Quantitative RT-PCR analysis demonstrated a near-complete absence of *Cfap47* mRNA transcription in the testis of *Cfap47*-KO mice (Fig. S30C). Although *Cfap47*-KO mice were healthy and exhibited normal development, *Cfap47*-KO male mice showed severely reduced fertility. The rate of two-cell embryos was markedly lower when using sperm from *Cfap47*-KO mice compared to wild-type (WT) mice (Fig. S30D). Despite similar sperm counts (Fig. S31E) and morphology (Fig. S31F) between *Cfap47*-KO and WT mice, sperm motility was significantly reduced in *Cfap47*-KO mice (Videos S2 & S3). Analysis of sperm motility revealed significant reductions in total and progressive motility in *Cfap47*-KO mice compared to WT mice (Fig. 7A). Notably, the percentages of non-linear and immotile sperm were significantly increased in the absence of CFAP47 (Fig. 7A). Furthermore, three kinematic parameters of sperm—curvilinear velocity (VCL), average path velocity (VAP), and straight-line velocity (VSL)—were significantly lower in *Cfap47*-KO mice than in WT mice (Fig. 7B). These results highlight the crucial role of CFAP47 in regulating sperm motility, supporting previous findings. For instance, earlier research discovered four patients with AZS carrying hemizygous *CFAP47* variants and described that *Cfap47*-KO mice are sterile with reduced sperm motility^43^. Further investigations identified more *Cfap47* mutations in AZS patients, emphasizing the critical role of CFAP47 in flagellar axoneme structure and its potential as a diagnostic target for AZS^44–46^.

**Figure 7.**
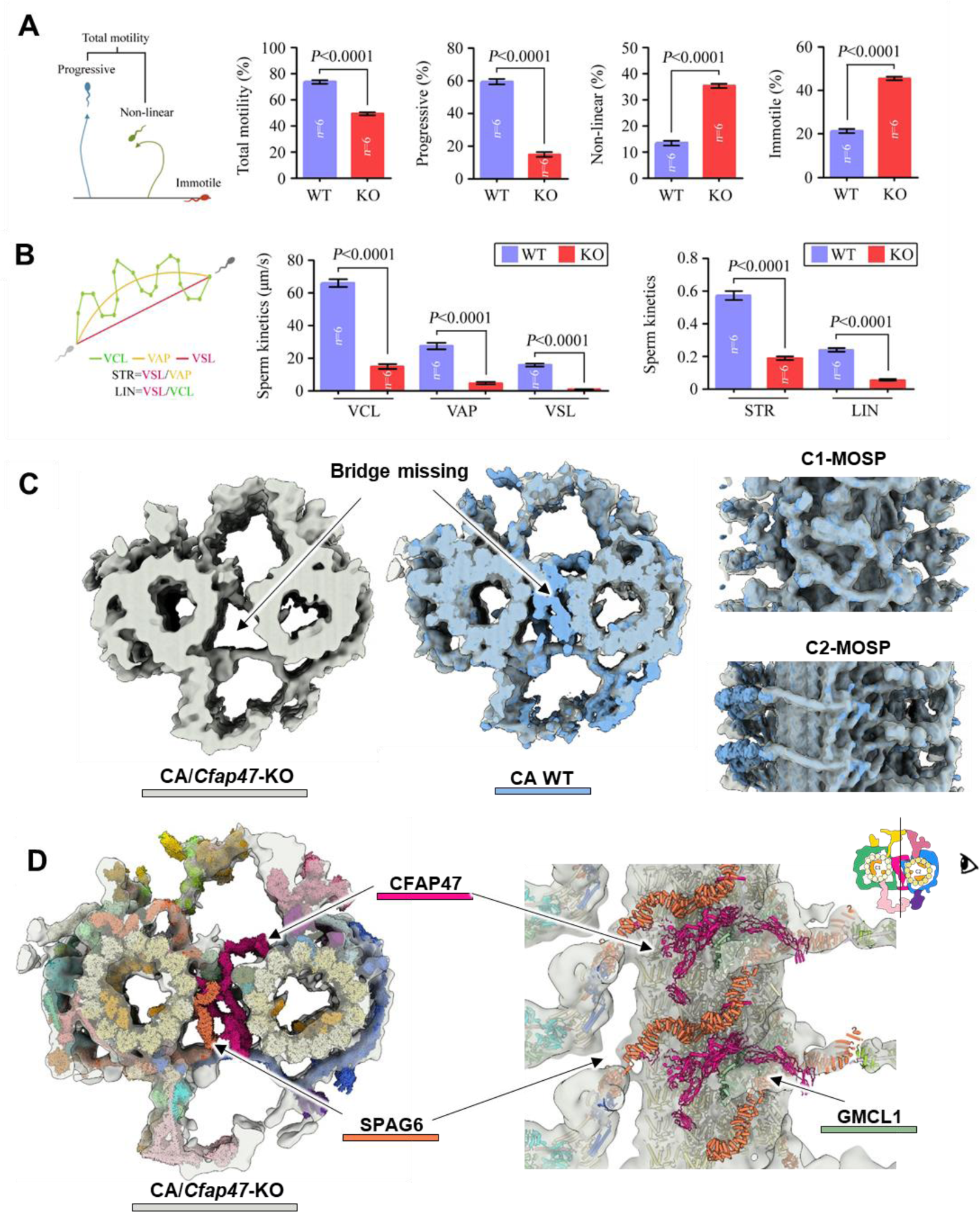
*Cfap47*-KO mice exhibit reduced progressive motility and impaired bridge density in CA structure. (**A**) Analysis of semen characteristics via a computer-assisted system, comparing total motility, progressive motility, non-linear motility, and immobility between WT and *Cfap47*-KO (KO) mice. (**B**) Measurement of curvilinear velocity (VCL), average-path velocity (VAP), and straight-line velocity (VSL) of sperm from WT and *Cfap47*-KO mice. Straightness (STR) is defined as VSL/VAP, and linearity (LIN) is defined as VSL/VCL. For all statistical analysis in this figure, student’s *t* test was used and error bars represented SD (*n*=6). (**C**) The overall structure of CA in *Cfap47*-KO mice (grey) is shown both individually and overlaid with WT CA structure (blue) in top view and side views. (**D**) The WT CA model fitted into the CA structure of *Cfap47*-KO mice, showing the top view and a side view from the bridge region. The *Cfap47*-KO mice exhibit absent density for CFAP47 and SPAG6 proteins in the bridge region of the CA structure.

Surprisingly, TEM analysis of sperm from *Cfap47*-KO mice showed an apparently normal “9+2” axoneme pattern^43^. Hence, the structural basis for CFAP47’s impact on sperm motility is quite perplexing. To investigate this further, we resolved the in situ structure of the sperm CA from *Cfap47*-KO mice. The raw tomogram revealed the “9+2” architecture of sperm axoneme in *Cfap47*-KO mice, yet the bridge region in CA is significantly more vacant compared to the WT CA (Fig. S30G). The sperm CA structure in *Cfap47*-KO mice was resolved to a resolution of 25 Å (Fig. S31A). Throughout the STA data processing, we observed greater structural heterogeneity in CA particles from *Cfap47*-KO mice compared to the WT. Ideally, 25% of CA particles picking at 8 nm intervals can contribute to the 32 nm repeat reconstructions. However, only about 11% (1k/9.3k) of the particles contributed to the 32 nm repeat CA structure in *Cfap47*-KO dataset (Fig. S31A), compared to around 18% (19k/105k) in the WT dataset (Fig. S2). This suggests that a significant portion of CA structures in *Cfap47*-KO sperm may be compromised.

The overall structure of the *Cfap47*-KO CA displays high similarity to the WT CA, except for the absence of density in most of the bridge region (Fig. 7C & Fig. S31B). By aligning the WT CA model with the mutant structure, several significant observations were made. Firstly, the entire CFAP47 density was absent, confirming the accuracy of the CFAP47 model identified in our CA structure (Fig. 7D). Secondly, the GMCL1 density remained intact in the mutant CA (Fig. 7D), indicating that although GMCL1 and CFAP47 closely interact to form the CA bridge, GMCL1 can exist independently of CFAP47 to link C1 and C2, thereby contributing to CA stability. Thirdly, the 16 nm repeating SPAG6 proteins that located on the bridge side of C1-MOSP, were absent in conjunction with CFAP47, suggesting that the assembly of SPAG6 in this region requires the presence of CFAP47 (Fig. 7D).

These results elucidate the structural mechanism underlying observed phenotype of reduced sperm motility in *Cfap47*-KO mice. On one hand, the mutant CA structure remains intact in C1/C2 MOSPs and all projections, allowing the CA to interact with RSs in all nine directions and thus enabling the assembly and movement of the 9+2 axonemal structure. This differs markedly from the absence of HYDIN, which affects the RS binding site structure on the half-circle. On the other hand, the absence of CFAP47 results in the hollowing of the CA central bridge, compromising the stability of the CA and potentially affecting the sustained movement and long-term structural stability of the whole axoneme. The comparison of *Cfap47*-KO and WT CA structures not only improved our comprehension of the structural and physiological roles of the CFAP47 protein, but also provided essential groundwork for understanding CFAP47-related human diseases.

## Discussion

The axoneme of cilia and flagella is a complex mechanical device that generates propulsion for swimming and fluid transport. Almost all motile cilia require CA as an indispensable core component, and the loss of the CA results in paralyzed cilia^12,47^. In this study, we employed the cutting-edge visual proteomics technology to obtain a significantly enhanced model of the extensively decorated microtubules in CA architecture, especially in higher mammals. The microtubule inner and outer surface proteins stabilize the CA and define its repeating units. The MOSP in the C1 and C2 microtubules wraps around the microtubule to serve as an assembly hub for all other projections. The chain-like proteins CFAP47 and HYDIN tethers the asymmetric C1 and C2 microtubules internally and externally in a complementary manner. These two longest ASH proteins ensure the orderly coordination of the C1 and C2 microtubules with different repeats, forming a stable CA structure that regulates the movement of the entire axoneme. Sperm from *Cfap47*-KO mice exhibited a hollowing of the CA central bridge, correlating with the phenotype of reduced sperm motility. These observations provide an essential basis for understanding the molecular assemblies of mammalian CA and their roles in regulating ciliary motility. Furthermore, our study exemplifies the integration of in-situ structural biology technology with gene-modified mouse models to explore defective cilia motility and ciliopathies at molecular level.

When aligning our structural model with previously reported map of human CA, we noticed that all protein structures, except for certain SPACA9 subunits in the C2 lumen, matched closely with the human CA structure (Fig. S32). It proves a high degree of structural and functional conservation of CA components between mice and humans. Consequently, the CA structures we have determined will be instrumental in guiding future studies on their mutations in human ciliopathies (Fig. S19). Mutations in these genes may alter residue types or cause protein truncations, potentially impacting CA function in at least two aspects. Firstly, mutations in MOSP and projections can directly impair the interaction between the CA and the corresponding RS heads. Secondly, mutations in scaffold proteins such as HYDIN and CFAP47 may impair the connection between the C1 and C2 microtubules, leading to an instability or even disassembly of the overall CA structure. Follow-up studies involving samples from patients with PCDs or mutant mouse models to explore more in situ structures of mutant CAs will provide more detailed insights.

Furthermore, our work presents a valuable approach for investigating the molecular mechanisms underlying CA-related sperm defects. Biomacromolecules such as CA are implicated in numerous human diseases, but their recombinant expression and purification in intact form are challenging. This difficulty hampers the study of disease-associated mutants at the molecular level. In this study, we successfully obtained the mutant CA structure from a modest dataset (95 tomograms, equivalent to only one day’s cryo-EM machine time) using a minimal amount of biological samples (less than 1% of a single mouse’s sperm). This enabled us to clearly observe the impact of *Cfap47*-KO on the overall CA structure and its influence on interacting proteins such as SPAG6, thereby elucidating the molecular mechanism by which CFAP47 affects sperm motility. Currently, the primary technology for analyzing patient with sperm defects is TEM analysis of resin-embedded sperm samples^48^. This method not only has lower resolution but also inevitably introduces sample artifacts or structural damage, potentially compromising the accuracy of analysis and interpretation. In contrast, the cryo-ET approach preserves samples in a near-native state, minimizes artifact formation, and achieves higher resolution structures. This method can be applied to analyze the in-situ structure of CA component proteins from individual patients, thereby uncovering the molecular-level pathogenesis of specific mutations.

According to our in situ reconstructions of the sperm CA, the complete axoneme structure is visible in the Z-axis slice (Fig. 1A). The nine RS heads are all closed to the projections and MOSPs on the CA. However, due to the highly dynamic nature of the RS and CA interface, we could not obtain the density map of these interface regions and thus cannot show the details of these interactions. The CA and RS has been proposed to regulate axonemal dynein activity through both mechanical and chemical signals^13^. Cryo-EM structures of RSs and their spoke heads have recently revealed that their distal surfaces are highly negatively charged^40^. However, the surface electrostatics of the distal surfaces of the C1a, C1b, C1c, and C2a projections, as well as the C1-MOSP and C2-MOSP, which are supposed to interact with the negatively charged surfaces of RS heads, mostly showed scattered regions of both positive and negative charges (Fig. S33). This suggests that RS and CA may not form a stable complex through the interaction of opposite surface charges. C1b and C2a predominantly showed highly negatively charged surfaces (Fig. S33), supporting a previously proposed model that electrostatic repulsion occurs between RS and CA when they are close enough during beating^40^. Detailed interactions between CA and RS require a more complete structural model of the CA-RS interface, including the flexible loop regions.

In a previously reported structure of *C. reinhardtii* CA^18^, it was proposed that the chain-like structure of CFAP47 within the bridge exhibited flexibility and elasticity, defining the extent of relative sliding and rotation between the C1 and C2 halves of the CA. Moreover, the C-terminal part of HYDIN was identified as the CA motor arm, occupying two different positions and states on the CA, suggesting its ability to slide along the microtubule track^18^. It was speculated that the movement of KLP1 arrays transmitted mechanical force from C2 to C1 through CFAP47, thereby affecting CA-RS interactions^18^. However, in our cryo-ET data, we did not observe significant sliding or rotations between the C1 and C2 halves. We also did not detect movement of the HYDIN-CTD along the C2 microtubule. The relatively stable connection between C1 and C2 in our structure is consistent with previously reported CA structures in other species (Fig. S5). Therefore, we speculate that there may be no significant movement between C1 and C2 in the axoneme, and the movement observed previously may be an artifact caused by the sample preparation method used for in vitro purification.

The application of AlphaFold2 has significantly advanced structural modeling based on secondary structural features in cryo-EM reconstructions at middle resolution^49^. This approach is applicable not only to protein complexes with known components, such as purified complexes, but also to the identification of previously unknown components of cellular complexes in their native enviroment^21^. It presents a powerful alternative to traditional genetic and cell biology methods for identifying protein components and localizing them within cells, without requiring labeling, cellular disruption, or purification, and is thus termed the visual proteomic approach^21^. In this study, we employed this cutting-edge technology to identify and construct pseudo-atomic models of 39 different proteins within the native mammalian CA structure. These proteins interact directly with the microtubule surface or are components of the microtubule-bound projections. Our findings significantly enhance our understanding of the proteins types that interact with microtubules and their mechanisms, with particular emphasis on the roles of the two longest ASH proteins, CFAP47 and HYDIN, which tethers the asymmetric C1 and C2 microtubules internally and externally in a complementary manner. Our identified proteins are highly conserved in humans and are closely associated with human ciliopathies. Consequently, our work provides a foundation for understanding CA-related human diseases, such as morphological abnormalities of sperm flagella (MMAF) and PCDs.

